# Biomass of a trophic level increases with maximum body size, but less than proportionally

**DOI:** 10.1101/2020.04.06.028381

**Authors:** Henrique C. Giacomini

**Affiliations:** Ontario Ministry of Natural Resources and Forestry, 2140 East Bank Drive, Peterborough, ON, Canada, K9J 7B8

## Abstract

A recent paper by Enquist and colleagues^1^ took a very important step in predicting the ecosystemic effects of species losses on a global scale. Using Metabolic Scaling Theory (MST), they concluded that large-sized species contribute disproportionately to several ecosystem functions. One of their key predictions is that total biomass of animals in a trophic level (*M*_*Tot*_, using their notation) should increase more than proportionally with its maximum body size (*m*_max_), following the relationship *M*_*Tot*_ ∝ *m*_max_^5/4^. Here I argue that this superlinear scaling results from an incorrect representation of the individual size distribution and that the exponent should be 1/4, implying a sublinear scaling. The same reasoning applies to total energy flux or metabolism *B*_*Tot*_, which should be invariant to maximum size according to the energetic equivalence and perfect compensatory responses entailed by MST.

---

The total biomass within a size interval characterizing a trophic level is calculated by integrating the product of individual mass *m* by its density function *f*(*m*), also known as size spectrum or individual size distribution:

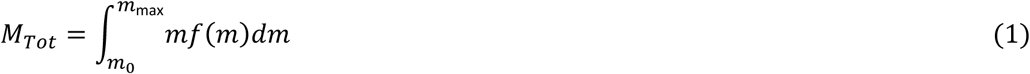

where *m*_0_ and *m*_*max*_ are the lower and upper limits of the size interval. To be an appropriate density function, *f*(*m*) must be measured as number of individuals per unit space (area or volume) per unit mass (ref. 2), so that, when integrated over a size interval, it gives the total number of individuals per unit space (denoted here by *N*_*Tot*_). This is also a necessary condition for *M*_*Tot*_ in equation (1) to be measured in the correct scale, in total mass per unit space.

Based on MST, the authors define the size spectrum of animals as a power function, i.e., *f*(*m*) = *cm*^−*ϵ*^, with *ϵ* = 3/4. However, the exponent −3/4 typically describes changes in abundance along logarithmic intervals of body size^3,4^, the so-called abundance spectrum^5^, in this case for a single trophic level (for multiple trophic levels the exponent is more negative due to energy losses from trophic transfers^6^). Such abundance already represents an integration over a size interval and is measured as number of individuals per unit space, so it is not suitable as a density funtion to calculate biomass in equation (1). The abundance spectrum of a trophic level can be represented as a function of the interval’s maximum size as *N*_*Tot*_ = *δm*_max_^−3/4^ (which reference size is used, whether maximum, minimum or a mid-point, does not change the exponent^7^). The relationship between *N*_*Tot*_ and *f*(*m*) can be thus expressed as:

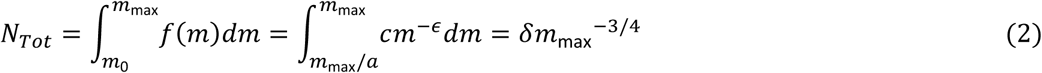

where *a* > 1 is a constant defining the ratio *m*_max_/*m*_0_, or the logarithmic range of the size interval; and *δ* is a coefficient given by *δ* = 4*c*(*a*^3/4^ − 1)/3. Most importantly, the exponent satisfying equation (2) is −*ϵ* = −7/4, one unit lower than the value used by Enquist et al. Similar demonstrations can be found in refs. 7-8.

The value of *ϵ* can be derived from more fundamental principles of MST if we assume a constant resource supply rate (*J*_*Tot*_, in mass per time per unit space)^3^. At equilibrium, the total metabolism or energy flux of a trophic level, *B*_*Tot*_, should be equal to their shared *J*_*Tot*_ (assuming further that *J*_*Tot*_ corresponds to assimilated energy), so that biomass remains constant in time:

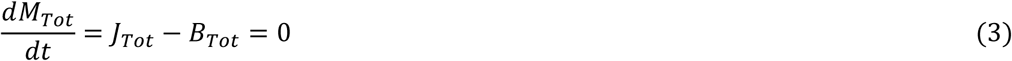

The total metabolism is equal to individual metabolism integrated over the size spectrum *f*(*m*). Individual metabolism is expected to scale as *∝ m*^3/4^ (ref. 3), so equation (3) can be expressed as:

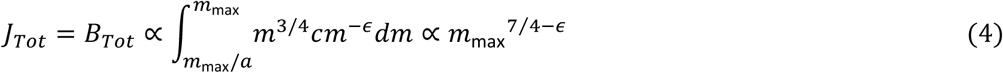

As *J*_*Tot*_ is invariant with respect to size (*J*_*Tot*_ ∝ *m*^0^), the equilibrium condition requires that *ϵ*= 7/4.

The assumption of constant resource supply underlies the energetic equivalence rule or hypothesis^4^ and explains the commonly observed scaling of *N*_*Tot*_ with an exponent -3/4 within a trophic level (or, more generally, with the reciprocal of individual metabolism’s). The reasons for energetic equivalence are still a debated topic, and the hypothesis have found mixed empirical support (e.g., refs. 9-10). It is nonetheless a prediction emerging from MST if all physiological processes determining the rates of energy gains and losses are proportional to individual metabolism and scale with the same exponent (e.g., 3/4). In this case, the efficiency with which energy is made available to biomass production, i.e., trophic transfer efficiency, can be represented by:

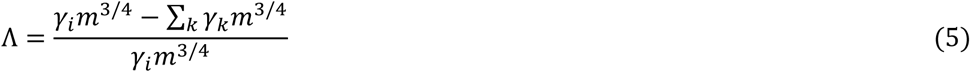

where *γ*_*i*_ is the coefficient for energy input (e.g., ingestion), and *γ*_*k*_ are the coefficients representing all processes leading to energy losses (i.e., not used by the next trophic level), including respiration, excretion, egestion, and mortality from causes other than predation. Given a common exponent, the body size component cancels out and efficiency becomes a constant: Λ = (*γ*_*i*_ − ∑_*k*_ *γ*_*k*_)/*y*_*i*_. Therefore, the amount of energy that is taken from a lower trophic level and metabolized remains the same (i.e., constant *J*_*Tot*_ and *B*_*Tot*_) regardless of which body sizes characterize the trophic level, and the resulting size spectrum scales as *f*(*m*) ∝ *m*^−7/4^ (equation 4), implying *N*_*Tot*_ ∝ *m*^−3/4^ (equation 2).

The total biomass calculated from equation (1) will be thus given by:

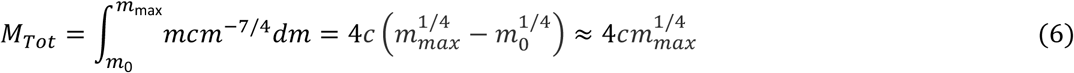

where the resulting power function is an approximation for small *m*_0_. It predicts an increase of biomass with maximum body size, but in a sublinear way.

It is important to recognize that the increase in biomass predicted by MST in equation (6) does not involve changes in ecological function. It results purely from the fact that larger organisms have slower turnover rates, which leads to greater biomass accumulation. From a community perspective, it implies perfect compensatory responses to species losses, as total resource utilization rate remains unchanged. Energy equivalence, constant trophic efficiency, and power-law size spectra are intrinsically related aspects of MST and all result from a common scaling of energetic processes. While there may be reasons for common scaling to result from long-term evolutionary processes^11^ or broad-scale macroecological averaging, many real cases of ecological interest involve transient short- or mid-term changes that deviate from MST assumptions. For instance, trophic interactions are generally size-structured, so smaller predators in a trophic level cannot immediately compensate for the loss of large ones. Due to the increased proportion of uneaten prey, trophic efficiency (equation 5) would decrease nonlinearly with size, preventing analytical solutions in the form of simple power functions and limiting the scope of MST. In those cases, MST predictions are not expected to match empirical data or simulations from more realistic models. One example is the 18% decline in total heterotrophic metabolism that resulted from megaherbivore losses in the General Ecosystem Model simulations by Enquist et al., in contrast with the MST expectation of no change. Nonetheless, MST still serves as a usefull baseline for comparison, one that controls for purely energetic processes but leaves out other important factors such as specialized ecological interactions and transient dynamics.

## References

1. Enquist, B.J., Abraham, A.J., Harfoot, M.B., Malhi, Y. & Doughty, C.E. The megabiota are disproportionately important for biosphere functioning. Nature Communications, 11(1) 1–11 (2020).

2. Andersen, K.H. & Beyer, J.E. Asymptotic size determines species abundance in the marine size spectrum. The American Naturalist, 168(1) 54–61 (2006).

3. Brown, J.H., Gillooly, J.F., Allen, A.P., Savage, V.M. & West, G.B. Toward a metabolic theory of ecology. Ecology, 85(7) 1771–1789 (2004).

4. White, E.P., Ernest, S.M., Kerkhoff, A.J. & Enquist, B.J. Relationships between body size and abundance in ecology. Trends in Ecology & Evolution, 22(6) 323–330 (2007).

5. Sprules, W.G. & Barth, L.E. Surfing the biomass size spectrum: some remarks on history, theory, and application. Can. J. Fish.Aquat. Sci. 73(4) 477–495 (2016).

6. Jennings, S. & Mackinson, S. Abundance–body mass relationships in size-structured food webs. Ecology Letters, 6(11) 971–974 (2003).

7. Blanco, J.M., Echevarría, F. & García, C.M. Dealing with size-spectra: some conceptual and mathematical problems. Sci. Mar. 58: 17–29 (1994).

8. Reuman, D.C., Mulder, C., Raffaelli, D. & Cohen, J.E. Three allometric relations of population density to body mass: theoretical integration and empirical tests in 149 food webs. Ecology Letters, 11(11) 1216–1228 (2008).

9. Isaac, N. J., Storch, D. & Carbone, C. Taxonomic variation in size-density relationships challenges the notion of energy equivalence. Biology Letters 7:615–618 (2011).

10. Hatton, I.A., Dobson, A.P., Storch, D., Galbraith, E.D. & Loreau, M. Linking scaling laws across eukaryotes. Proceedings of the National Academy of Sciences, 116(43) 21616–21622 (2019).

11. Hin, V. & de Roos, A.M. Evolution of size-dependent intraspecific competition predicts body size scaling of metabolic rate. Functional Ecology, 33(3), 479–490 (2019).

